# The mobilizable plasmid P3 of *Salmonella enterica* serovar Typhimurium SL1344 depends on the P2 plasmid for conjugative transfer into a broad range of bacteria *in vitro* and *in vivo*

**DOI:** 10.1101/2022.09.15.508199

**Authors:** Marla Sofie Gaissmaier, Leanid Laganenka, Mathias Klaus-Maria Herzog, Erik Bakkeren, Wolf-Dietrich Hardt

**Affiliations:** Institute of Microbiology, ETH Zürich, Switzerland; Department of Zoology and Department of Biochemistry, University of Oxford, Oxford, UK

## Abstract

The global rise of drug-resistant bacteria is of great concern. Conjugative transfer of antibiotic resistance plasmids contributes to this antibiotic resistance crisis. Despite the substantial progress in understanding the molecular basis of conjugation *in vitro*, the *in vivo* dynamics of intra- and interspecies conjugative plasmid transfer are much less understood. In this study, we focused on the streptomycin resistance-encoding mobilizable plasmid pRSF1010^SL1344^ (P3) of *Salmonella enterica* serovar Typhimurium (*S*. Tm) strain SL1344. We show that P3 is mobilized by interacting with the conjugation machinery of a second, conjugative plasmid pCol1B9^SL1344^ (P2) of SL1344. Thereby, P3 can be transferred into a broad range of relevant environmental and clinical bacterial isolates *in vitro* and *in vivo*. Our data suggests that *S*. Tm persisters in host tissues can serve as P3 reservoirs and foster transfer of both, P2 and P3 once they reseed the gut lumen. This adds to our understanding of resistance plasmid transfer in ecologically relevant niches including the mammalian gut.

**IMPORTANCE:** *S*. Tm is a remarkably adaptable and globally abundant bacterial species that rapidly occupies new niches and survives unstable environmental conditions. As an enteric pathogen, it can potentially interact with a broad range of bacterial species residing in the mammalian gut. High abundance of bacteria in the gut lumen facilitate conjugation and spread of plasmid-encoded antibiotic resistance genes. By studying the transfer dynamics of the P3 plasmid *in vitro* and *in vivo*, we illustrate the impact of *S*. Tm-mediated antibiotic resistance spread via conjugation to a variety of relevant environmental and clinical bacterial isolates. Along with temperate phages or naked DNA, plasmids are among the most critical vehicles driving antibiotic resistance spread. Further understanding of the dynamics and drivers of antibiotic resistance transfer, along with identifying the environmental niches where this occurs, is needed to develop effective solutions for slowing down the emerging threat of multidrug-resistant bacterial pathogens.

## INTRODUCTION

Bacterial infections pose a high risk to human health. The use and over prescription of antibiotic treatments in human and veterinary medicine has been linked to the increasing emergence of antibiotic-resistant bacteria (1–3). According to the priority list published by the WHO concerning the increase of multidrug resistant bacterial species, members of the *Enterobacteriaceae* family were classified as “most critical” (4). Further, recent reports published by *The Lancet* claim that almost 5 million deaths were associated with antibiotic resistance in 2019 (5). Similarly, the spread of animal- or plant-associated pathogens with antibiotic resistance causes high treatment costs in agriculture and reduces the yield of certain crops (3, 6, 7). Understanding the mechanisms and slowing down the spread of antibiotic resistances of opportunistic or pathogenic bacteria is thus an important task to preserve antibiotics as an effective treatment strategy.

Horizontal gene transfer (HGT) is a dominant mechanism for acquiring antibiotic resistance (8). HGT mostly occurs by means of transformation, transduction or conjugation (9). Out of these mechanisms, conjugation of plasmids is arguably the most important driver for spreading antibiotic resistance genes amongst bacteria (10–12). Plasmids are usually circular, self-replicating DNA molecules. In contrast to chromosomal DNA, they do not encode essential house-keeping genes, but rather accessory genes that help their host to adapt to particular environments and are linked to new metabolic functions, antibiotic resistance or virulence (8, 13, 14). Generally, plasmids can be classified into three categories based on their mobility: 1) conjugative plasmids that encode their own conjugation machinery, 2) mobilizable that do not encode a functional conjugation machinery, and 3) non-mobilizable plasmids. It was estimated that conjugative and mobilizable plasmids make up around one-half of described plasmids (15).

In this study we focused on the transfer dynamics of the mobilizable plasmid pRSF1010^SL1344^ (termed P3, Fig S1) which naturally resides in strain *S.* Tm SL1344, an isolate from cattle that belongs to a clade of *Salmonella* strains contributing significantly to infections world-wide (16–18). P3 belongs to the IncQ incompatibility family of plasmids and is a close relative of the RSF1010 plasmid originally isolated from *Escherichia coli* (17, 19, 20). Similar to P3, it carries streptomycin *(strAB)* and sulfonamide resistance (*sulII*) genes that are typically encoded within the same gene cluster and occur in various bacterial species (17, 21). Furthermore, previous studies showed that IncQ plasmids were capable of taking up genes conferring resistance to other clinically relevant antibiotics like gentamycin or kanamycin (22). *S*. Tm strain SL1344 additionally harbors two conjugative plasmids pSLT^SL1344^ and pCol1B9^SL1344^, which will be referred to as P1 and P2, respectively (23). *S*. Tm typically contains 12 - 15 copies of P3, whereas only 1 to 2 copies of P1 and P2 are present per cell (19, 24, 25). In contrast to P3, which only harbors an origin of transfer *(oriT)* and genes required for mobilization, the plasmids P1 and P2 encode complete conjugation machineries (19, 26, 27). P3 also encodes a plasmid-derived replication machinery (helicase, primase and iteron-specific DNA-binding protein), allowing it to replicate in a host-independent manner (19, 28). Notably, the high similarity to the broad host range plasmid RSF1010 suggests that P3 might have a broad host range as well (Fig S1b). For our study, this was of particular interest in combination with a host bacterium like *S*. Tm that can grow both inside and outside of the intestinal tract of host animals. Furthermore, this pathogen can form persister reservoirs (that is subpopulations of bacteria that survive upon exposure to antibiotics) in host tissues that might “store” plasmids over long periods of time (23, 29). Importantly, the persister populations can migrate back to the gut lumen after the antibiotic treatment has ended and resume gut-luminal growth (re-seeding), further interacting with a significant number of bacterial species which inhabit or pass through the animal gut (30–32).

We show that P3 of *S.* Tm can be mobilized by employing the conjugation machinery of the co-occurring P2 plasmid. Moreover, we found evidence for conjugational transfer of P3 to a variety of Gammaproteobacteria members not only *in vitro* but also in the animal gut. As *S*. Tm persisters residing in host tissues can serve as P3 reservoirs and foster conjugative transfer of both, P2 and P3 once they reseed the gut-lumen, we speculate that the animal gut may be a relevant niche for spreading P3.

## RESULTS

### P3 requires P2 for conjugational transfer into a broad host range *in vitro*

As P3 is transferred to other *S*. Tm strains (Fig. 1a) but lacks the genes for a conjugation apparatus, we asked if the conjugative plasmids P1 or P2 of *S*. Tm SL1344 might facilitate conjugative P3 transfer. *S*. Tm ATCC 14028S, which naturally lacks the plasmids P2 and P3, was used as a recipient strain and transconjugants were detected after 2 h mating between SL1344 and 14028S in liquid medium (Fig 1a). However, no transconjugants were observed when the donor strain is cured of P2. This suggests that P2 is required for transfer of the P3 plasmid. Disruption of the P3 origin of transfer, *oriT*, similarly abolished plasmid transfer, further confirming that it is indeed conjugation that drives the transfer of P3 (Fig 1a).

**Figure 1:**
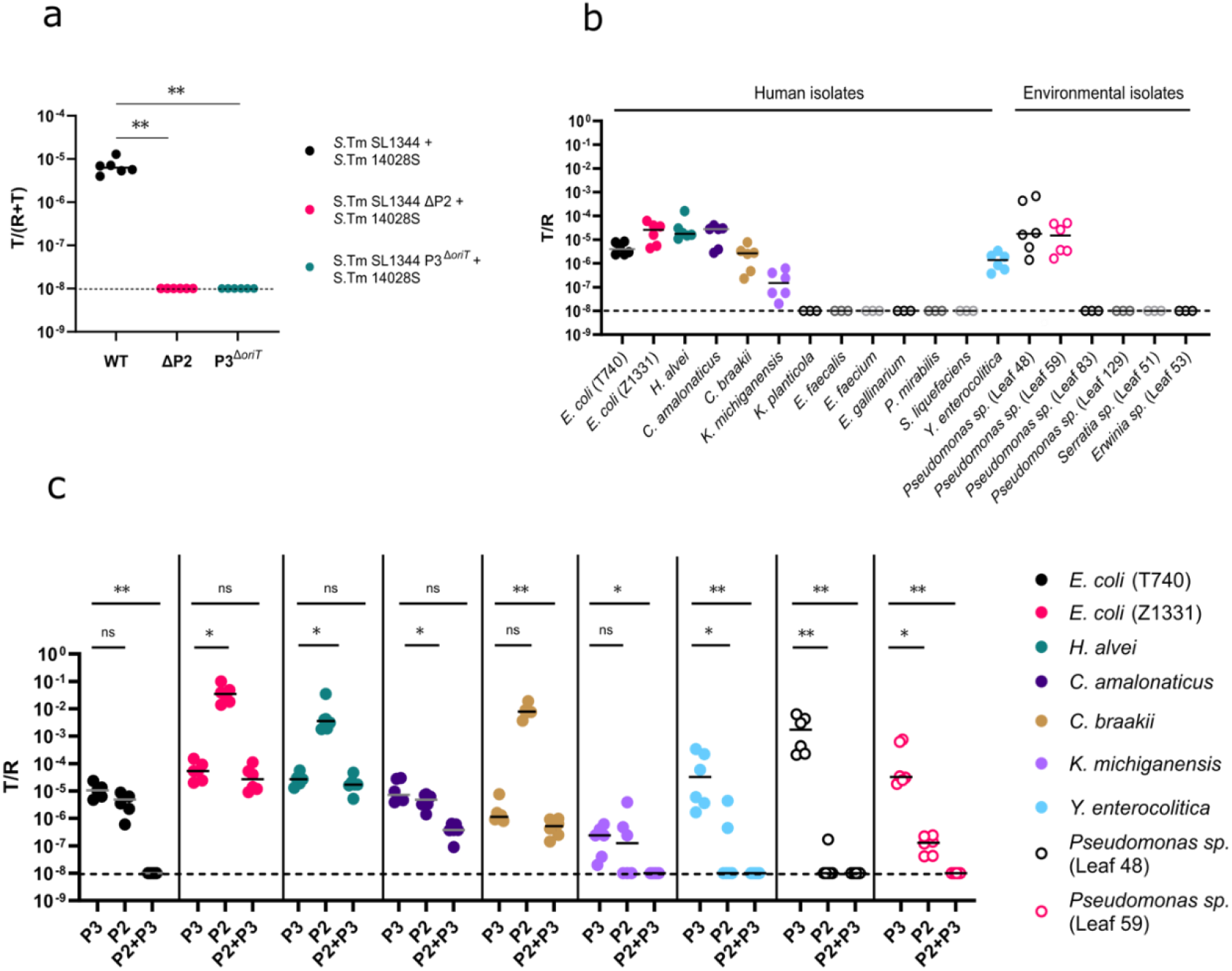
P3 is mobilized by P2 and can be stably maintained in a broad range of bacteria. **a | P2 is required for P3 transfer.** Final conjugation frequencies (T/R+T) of P3 for 2 h liquid mating of the three depicted donor/recipient combinations are shown. *S.* Tm SL1344 and *S.* Tm 14028S recipient were mixed 1:1 in LB and incubated for 2 h. Respective plating on MacConkey plates was used for enumeration of donor, recipient and transconjugant counts. n = 6 for each group; two independent experiments, p = 0.0022, two-tailed Mann-Whitney test. LOD = 10^-8^ **b | *In vitro* host range of P3.** The ratio of transconjugants to recipients after overnight agar mating. 1: 1 mix of the depicted recipients and S. Tm SL1344 donor was incubated on LB agar overnight. n = 3 - 6 per group, LOD = 10^-8^ **c | Stable maintenance of P2 and P3 depends on the recipient strain.** Ratio (T/R) of P3, P2 and P2+P3 transconjugants after overnight agar mating. 1:1 mix of recipient + S. Tm 14028S P2^*cat*^ P3 was incubated on LB agar overnight. n= 6 per group, two independent experiments. Kruskal-Wallis Test, comparing P3 with P2 or P3 with P2+P3, LOD = 10^-8^, p-value style: 0.1 (ns), 0.01(*), 0.001 (**).

To study the host range of the P3 plasmid, environmental and human bacterial isolates were selected based on the spectrum of antibiotic resistance (which should be different than that of *S.* Tm SL1344 donor strain) and ability to ferment lactose. This enabled detection and enumeration of P3 transconjugants based on differential plating or morphological differentiation of lactose-negative donor and lactose-positive recipient strains on MacConkey agar plates (Fig S2). The host range of P3 was investigated by performing overnight liquid and surface mating. For liquid mating, transconjugants could be detected in several human (*Escherichia coli, Hafnia alvei, Citrobacter amalonaticus, Yersinia enterocolitica*) and environmental isolates (*Pseudomonas* sp. Leaf48 and Leaf59), albeit with different transconjugant yields (Fig S3). PCR was used to verify P3 in the transconjugants (Fig S4a). In contrast to previous work on other RSF1010 plasmids showing conjugative transfer to Gram-positive bacteria (33), no transconjugants were detected for Gram-positive representatives of the *Enterococcaceae* family (*Enterococcus faecalis, E. faecium, E. gallinarium*) (Fig 1b, S3). We speculate, that this might be related to the specificity of the P2 sex pilus. Surface mating additionally enabled conjugative transfer of P3 into *C. braakii* and *Klebsiella michiganensis*, likely due to a closer contact between bacteria grown on a surface of an agar plate rather than in the liquid medium (Fig 1b). The average ratio of detected CFUs for transconjugants to recipients (ration_(T/R)_) remained in a range of 10^-6^ – 10^-4^ for most tested strains, similar to the observed ratios in liquid mating. Although still above the detection limit, the ratio(T/R) of transconjugants for *K. michiganensis* was lower at around 5 x 10^-8^.

All strains that showed transconjugants were consequently mated with *S*. Tm SL1344 *P3^ΔoriT^* to verify that plasmid uptake indeed occurred via conjugation and not by any other mechanism of horizontal gene transfer. No streptomycin-resistant CFUs indicative of transconjugants were detected for any of the tested recipient strains when using an *oriT* deficient donor, suggesting that conjugation was indeed the sole mechanism responsible for P3 transfer in all recipient strains (Fig S4b).

### P2 and P3 are often co-transferred into the same recipients

As our data above showed that P2 is required for P3 transfer, it was interesting to compare the P2 host range and conjugation rate to that of P3. We chose *S*. Tm 14028S harboring P2^*cat*^ and P3 as donor for all mating pairs, allowing us to follow the dynamics of P2 transfer by differential plating using chloramphenicol as a selection marker. After overnight agar mating, P2^*cat*^ was detected in high numbers for the tested *E. coli* and *H. alvei* strains. Both tested environmental samples *(Pseudomonas* sp. Leaf48 and Leaf59) as well as *K. michiganensis* and *Y. enterocolitica* showed low or undetectable levels of P2^*cat*^ transconjugants (Fig 1c). These findings suggest that although P3 stably replicates upon transfer into a recipient strain, P2 seems to be mostly lost from the *Pseudomonas* sp. Leaf48 and Leaf59, *K. michiganensis* and *Y. enterocolitica* recipient cells within 24 h. With P2 being a large and low copy plasmid, possible reasons for plasmid loss could be high fitness burden on the host or segregational loss (34). We additionally analyzed the genomes of the recipient strains for plasmids of the same incompatibility complex as that of P2 (Incl1). No potentially incompatible plasmids were detected in the strains based on our BLAST analysis of the conserved replication initiation protein RepZ of IncI1 plasmids (35). Interestingly, subsequent surface mating of all P3 transconjugants with a naïve recipient *E. coli* strain (*E. coli* W3110 *tsr::kmR)* further confirmed that conjugative P3 transfer was only possible for donor cells inheriting both P2 and P3 plasmids (Fig S4c).

### P3 transfer *in vivo*

The main ecological niche occupied by the P3 plasmid carrying *S*. Tm is the mammalian gut. We therefore used a well-established mouse colitis model (23, 27, 36) to verify the previous *in vitro* results and to study the dynamics of P3 transfer in this niche. C57BL/6 specific pathogen free (SPF) mice were pretreated with ampicillin to suppress the microbiota and allow the strains of interest to grow up to high densities, which was necessary for successful conjugation (27). The mice were then infected with respective recipient strains 24 h prior to infection with the *S*. Tm SL1344 donor strain. The number of transconjugants was determined by daily collecting and differential plating of feces and cecal content for the next 3 days. Transconjugants were detected in *E. coli* Z1331, *H. alvei*, *C. braakii*, *Y. enterocolitica* and *Pseudomonas* sp. Leaf59 (Fig 2a-e, respectively). No transconjugants were detected in *K. michiganensis* (Fig 2f). This shows that P3 indeed gets transferred between species within the gut system if donor and/or recipient strains can grow up to high densities.

**Figure 2:**
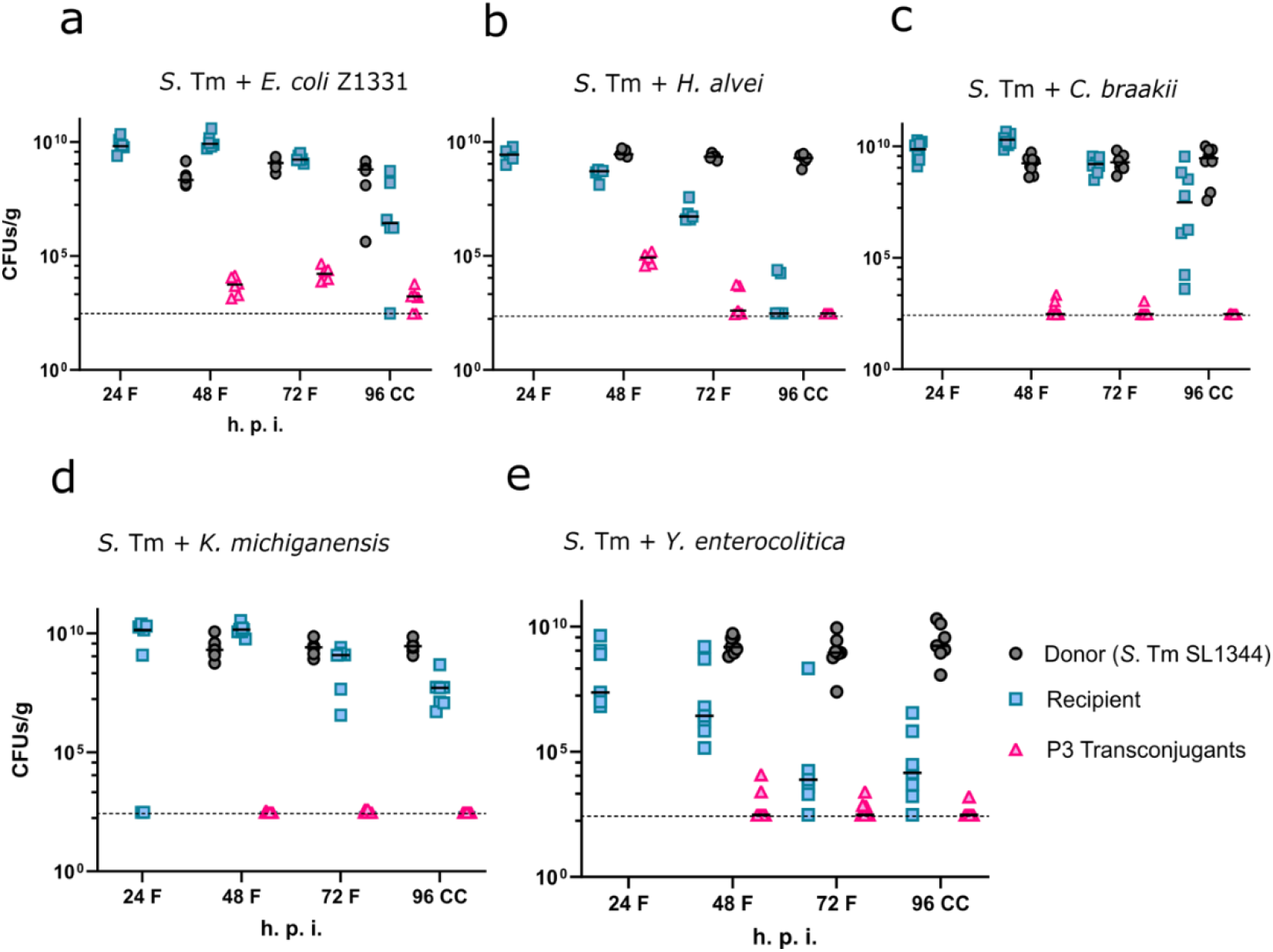
P3 transfer occurs in the gut and is not limited to one species. Total CFU count of donor, recipient and transconjugant populations in the feces and cecal contents. Ampicillin-pretreated C57BL/6 mice were infected with 5 x 10^7^ CFUs of the recipient strain on day 0 and with 5 x 10^7^ CFUs of the donor strain *S*. Tm SL1344 on day 1. Donors, recipients and transconjugants are indicated as grey circles, cyan squares and pink triangles, respectively. Tested recipients are **a |** *E. coli* Z1331 WITS1-AmpR, lac^+^, n = 6; **b |** *H. alvei* pM965 (Amp^R^) pACYC184 (Cm^R^), n = 5; **c |** *K. michiganensis* (Amp^R^, lac^+^), n = 4; **d |** *C. braakii* pM965 (Amp^R^, lac^+^), n = 8; **e |** *Y. enterocolitica* pM965 (Amp^R^, Km^R^), n = 4; **f |** *Pseudomonas* sp. Leaf 59 (Amp^R^, Cm^R^), n = 3.

### Reseeding *S*. Tm persisters as P3 reservoir

*S*. Tm can invade not only the gut tissue, but also colonizes systemic organs and forms long-term reservoirs at these sites (37–39). From these reservoirs, the pathogen can again pass the epithelial barrier and thereby reseed into the gut lumen. Specified pathogen free *Nramp^+^* 129S6/SvEvTac mice were used to assess the contribution of donor tissue reservoirs and reseeding events to the spread of P3 in the gut lumen. This mouse line is more resistant to *S.* Tm infection and thus more suitable for long term infections than the C57BL/6 mice used above. On the first day, mice were intraperitoneally (i.p.) infected with the SL1344 donor strain (*S.* Tm SL1344 P3^SmR^ pM965^ApR^). I.p. infections enabled direct infection of the systemic sites while bypassing the gut system. This was necessary to maintain a donor-free gut lumen and establish systemic *S.* Tm infection at the same time. Two days post infection (d.p.i.), the mice were orally treated with ampicillin (to suppress the microbiota and thereby promote gut luminal recipient growth) and subsequently infected with the 14028S recipient strain (*S.* Tm ATCC 14028S *marT::cat^CmR^)* by oral gavage. The dynamics of donor reseeding and subsequent conjugation were quantified for the following 6 days by collecting and differential plating of feces and cecal content (Fig 3a, b). Donor reseeding occurred typically within the first 3 days post antibiotic treatment, and the donor population was able to grow up to the carrying capacity (10^9^ CFUs/g feces) within one day after reseeding (Fig 3c, Fig. S6a). P3 transconjugants were detected in all tested mice as soon as reseeding had occurred. No transconjugants could be detected when *S*. Tm SL1344 P3^Δ*oriT*^ was used as plasmid donor, further confirming that conjugation is the only route for P3 transfer in this model (Fig S5).

**Figure 3:**
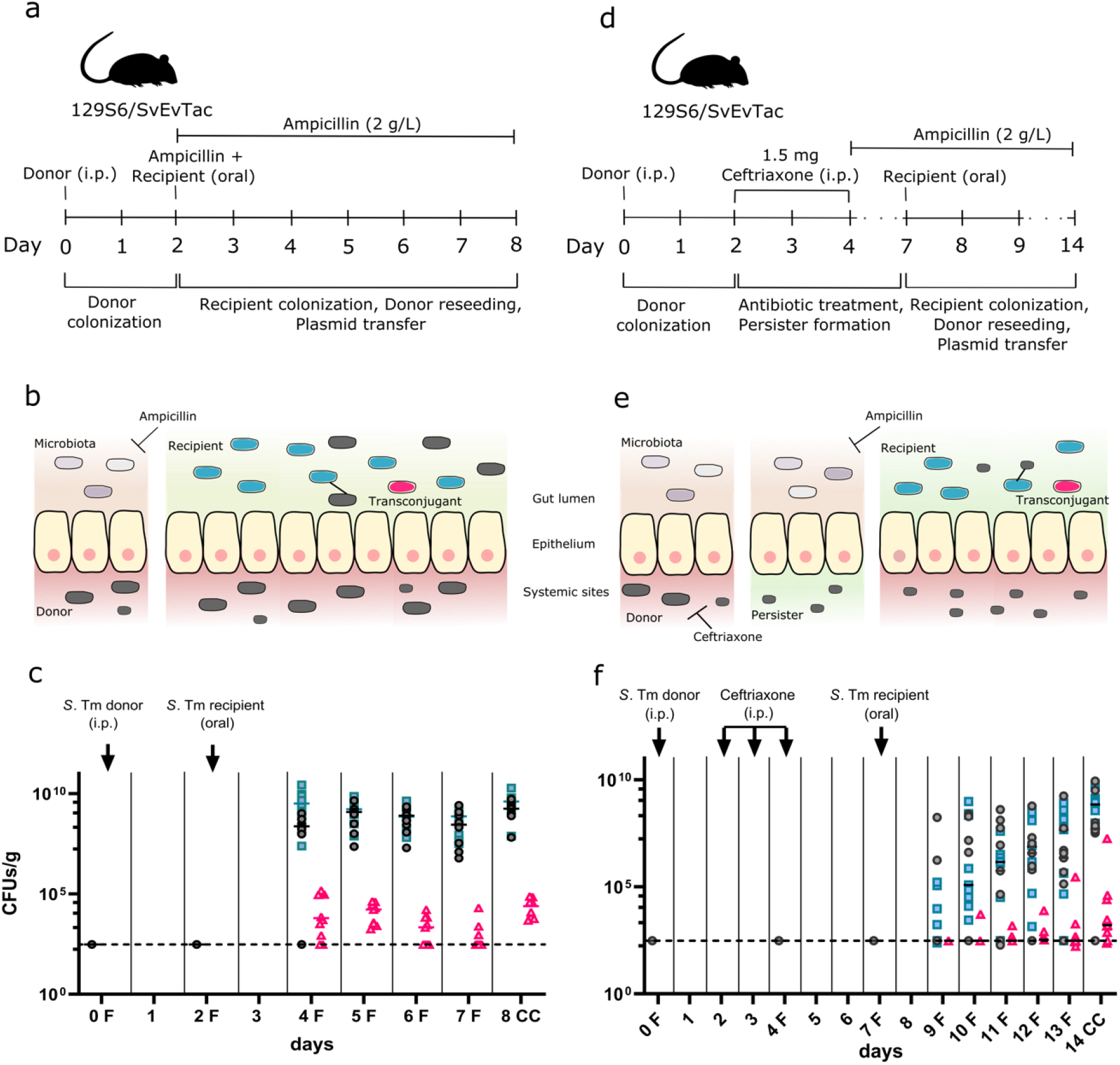
Systemic infection with *S.* Tm SL1344 shows conjugative P3 transfer after donor reseeding from tissue reservoirs into the gut lumen regardless of antibiotic treatment. **a | Experimental setup for reseeding donor model adapted from Bakkeren et al. (23, 29)** The reseeding donor model contained two phases: 1.) donor colonization and establishment of tissue reservoirs at systemic sites. 2.) recipient colonization in the gut lumen, followed by donor reseeding into the lumen and subsequent plasmid transfer via conjugation. **b | Reseeding model.** *S*. Tm donors (grey) establish tissue reservoirs after intraperitoneal injections and spread to organs. The microbiota provides colonization resistance against colonization of the gut lumen. Recipients (cyan) colonize the gut lumen by an oral infection following an ampicillin treatment to suppress the microbiota (presence of antibiotics is indicated by green shading). *S*. Tm donors reseed from their reservoirs and transfer plasmids to the recipient (indicated as line) and forming transconjugants (pink). **c | Conjugative P3 transfer by reseeding donors from tissue reservoirs into the gut lumen.** Mice were infected with 5 x 10^7^ CFUs of the donor *S.* Tm SL1344 (Amp^R^, Km^R^,Sm^R^) on day 0 (i.p.) and with 5 x 10^7^ CFUs of the recipient *S.* Tm 14028S (Amp^R^, Cm^R^) on day 2 (oral), 4 h post oral ampicillin pretreatment (20 mg). 2 g/L ampicillin was additionally added to the drinking water on day 2 post infection and maintained throughout the experiment. CFU counts of donor, recipient and transconjugant populations in the feces and cecal contents were determined by differential plating. Two mice were excluded from the analysis as they developed severe symptoms and were euthanized earlier for ethical reasons. *S.* Tm was invariably detected in the systemic sites (Fig S6a). Donors, recipients and transconjugants are indicated as grey circles, cyan squares and pink triangles, respectively. F: fecal sample, CC: cecal content. n = 8, two independent experiments. **d | Experimental setup for reseeding persister model adapted from Bakkeren et al. (23, 29)** The reseeding persister model contained three phases: 1.) donor colonization and establishment of tissue reservoirs at systemic sites. 2.) clearance with antibiotics and persister formation at systemic sites 3.) recipient colonization in the gut lumen, followed by donor reseeding into the lumen and subsequent plasmid transfer via conjugation **e | Persister model.** *S*. Tm donors (grey) establish tissue reservoirs after intraperitoneal injections and spread to organs. The microbiota provides colonization resistance against colonization of the gut lumen. Upon ceftriaxone treatment only persisting *S*. Tm cells can survive which leads to new tissue reservoir formation (presence of antibiotics is indicated by green shading). Recipients (cyan) colonize the gut lumen enabled by oral infection following an ampicillin treatment (supplemented in the water) to suppress the microbiota. *S*. Tm donors reseed from their newly established reservoirs, transfer plasmids to the recipient and thereby forming transconjugants (pink) (23). **f | Conjugative P3 transfer by reseeding persisters from tissue reservoirs into the gut lumen.** Mice were infected with 5 x 10^7^ CFUs of the donor *S.* Tm SL1344 (Amp^R^, Km^R^, Sm^R^) on day 0 (i.p.) and with 5 x 10^7^ CFUs of the recipient *S*. Tm 14028S (Amp^R^, Cm^R^) on day 7 (oral). 2 g/L ampicillin was added to the drinking water on day 4 and maintained throughout the experiment to suppress the gut microbiota for subsequent oral infection with the recipient strain. CFU counts of donor, recipient and transconjugant populations in the feces and cecal contents were determined by differential plating. One male mouse had to be excluded from the analysis as he developed severe symptoms on day 10 and was euthanized for ethical reasons. *S.* Tm was invariably detected in the systemic sites (Fig S6b). Donors, recipients and transconjugants are indicated as grey circles, cyan squares and pink triangles, respectively. F: fecal sample, CC: cecal content. n = 9, two independent experiments.

Finally, we assessed the transfer dynamics of P3 for persisting *S.* Tm donors. Antibiotic persistence describes the survival of a small subpopulation of bacteria upon exposure to antibiotics (40–42). In contrast to genetically inherited tolerance or resistance which affect the whole population, a persisting subpopulation will survive and re-establish reservoirs by clonal expansion after antibiotic treatment but stays sensitive to the respective antibiotic (43). Furthermore, studies revealed that *S*. Tm persisters residing at systemic sites can subsequently re-enter the gut lumen (44) and can serve as P2 reservoir (23). These tissue-lodged persister reservoirs are of special concern as they survive antibiotic treatments for a longer period of time and can lead to recurring or chronic infections (45). Because our previous *in vitro* experiments implied that P2 can act as a helper plasmid for mobilizing P3, we did a follow-up experiment to detect whether P3 would be transferred along with P2 if both derive from persistent *S*. Tm donors. To focus our analysis on the plasmids which are carried by tissue-lodged persisters, we extended the reseeding model described above by a 3-day i.p. ceftriaxone treatment starting 2 days post donor infection. This would eliminate all *S*. Tm cells growing within the mouse, while tissue-lodged persisters would prevail. The drinking water was supplemented with ampicillin (2 g/L) to suppress the gut microbiota and open a niche for the recipient (Fig 3d, e). The 14028S recipient strain was orally introduced on day 7. Again, the dynamics of donor reseeding and subsequent conjugation were quantified for the following 7 days by collecting and differential plating of feces and cecal content. Persister reseeding as well as P3 transfer to the recipient strain was detected in 7 out of 9 mice by 14 d.p.i. (Fig 3f, Fig. S6b). The lower counts of P3 transconjugants in this experimental setup (Fig. 3d-f) compared to the reseeding model (Fig. 3a-c) could be attributed to lower gut-luminal densities of both donor and recipient strains and higher variation in CFUs between the animals. Nevertheless, our data clearly shows that *S*. Tm persisters serve as antibiotic resistance plasmid reservoir which leads to subsequent gut-luminal transfer not only of conjugative plasmids such as P2 as shown in previous studies (23) but also of mobilizable plasmids such as P3, as shown here.

## DISCUSSION

Conjugative plasmid transfer is a critical driver of antibiotic resistance spread among bacteria (10–12). Investigating the distribution and transfer dynamics of broad host range plasmids is of particular importance, as they can potentially contribute to HGT between distantly related bacterial species. This study focused on the host range and transfer dynamics of the mobilizable plasmid P3 of *S*. Tm SL1344 *in vitro* and *in vivo*, which is part of the broad host range IncQ plasmid family. As transfer of mobilizable plasmids is dependent on external genes that encode the conjugation machinery, we aimed at determining the source of the conjugation machinery explaining P3 transfer. Our data suggests that the conjugative P2 plasmid of *S*. Tm SL1344 is responsible for P3 transmission. P3 transfer was additionally abolished by deletion of its origin of transfer (*oriT*), further confirming that it is indeed conjugation and no other type of HGT that drives P3 transfer. Since we have not tested a P1-deficient *S*. Tm donor strain, no final assumptions about its importance for P3 mobilization can be made. However, it is known that P1 is less efficiently transferred via HGT than P2 and SL1344 donors only harboring P1 were not able to transfer P3 into the recipient 14028S strain (26, 46).

Because they can replicate in a variety of bacteria, broad host range plasmids like members of the mobilizable IncQ-family are of special concern if they harbor antibiotic resistance genes. Plasmids of the IncQ family are usually associated with streptomycin and sulfonamide resistance but additional clinically relevant antibiotic resistance genes have been reported (19, 22). Hence, it was interesting to further explore the P3 host range by testing clinically and environmentally relevant isolates of the Proteobacteria phylum, as multiple members are of special concern in the emergence of antimicrobial resistance spread according to WHO. Therefore, a panel of potential recipients, mainly Gram-negative *Enterobacteriaceae* members that would encounter *S*. Tm in a natural environment like the human or animal intestine, was screened. Additionally, environmental *Pseudomonas* and *Erwinia* isolates were tested as potential P3 recipients (47), as streptomycin used to be an important agricultural bactericide and emerging streptomycin-resistant plant pathogens are of serious concern e.g. for fighting fire blight in orchards (3). Overnight *in vitro* liquid and surface mating with *S*. Tm SL1344 donors showed that P3 can indeed be transferred to several relevant microbiota members. Transconjugants were detected for the following human isolates: *E. coli* Z1331, *E. coli* T740, *H. alvei, K. michiganensis, C. amalonaticus, C. braakii* and *Y. enterocolitica*. These findings are in line with previous studies that were able to show the broad host range of RSF1010-like plasmids (19, 48, 49). Our study extends the reported list by including *H. alvei*, *Y. enterocolitica* and *Citrobacter* sp.. Interestingly, P3 transconjugants were also detected in two environmental isolates of *Pseudomonas* sp. (Leaf48 and Leaf59). Although the genes conferring streptomycin resistance (*strAB*) originally resided on transposons, their transfer by broad-host-range plasmids can potentially contribute to a rapid spread of streptomycin resistance in both commensal and pathogenic plant-associated bacteria (21). Nevertheless, our study implies that the spread of P3 in a given population is limited by the efficiency of P2 transfer and maintenance within the recipient cell. We were able to show that emergence of P3 transconjugants does not necessarily coincide with successful P2 plasmid transfer or maintenance. However, if the recipient cell does not harbor another plasmid that could be exploited by P3 for further conjugation, no further spread of antibiotic resistance by HGT is possible. The reasons behind the differential ability of transconjugants to stably maintain the P2 plasmid are yet to be studied. Possible explanations include segregational loss due to low copy numbers and high fitness costs connected to the maintenance of the plasmid (34, 50, 51).

In conclusion, our *in vitro* experiments show the dynamics of P3 plasmid transfer and how a mobilizable plasmid like P3 can take advantage of conjugative plasmids to enter new host cells. Compared to P2, which is lost rapidly by most recipients, P3 could have some advantageous traits such as expressing a host independent replication machinery, a small size which might impose little metabolic burden upon its host, and by a higher copy number of around 12-15 per cell (19). Further experiments should be conducted to properly assess the stability of P3 and its impact on host fitness over longer periods of time to assess how stable it might be maintained in environmental niches.

Our in vivo studies verify that the animal gut is a niche permitting resistance plasmid transfer. The data extend previous work by showing P3 transfer into *H. alvei* recipients, *C. braakii*, *Y. enterocolitica* and *Pseudomonas* sp. Leaf59.

On top, we established that tissue resident *S*. Tm cells can serve as P3 donors after re-seeding the gut lumen. Again, this transfer occurred independently of selective pressure for the streptomycin resistance encoded on P3 and the P3 transconjugant population was stably maintained at a level of up to 10^5^ CFUs/g feces or cecal content for reseeding donors throughout the experiment.

Persisters residing at systemic sites are of special concern as they can survive antibiotic treatments over long periods of time and can lead to reoccurring infections by reseeding into the gut lumen (52). Our data extends this previous work by demonstrating P3 transfer from persister reservoirs. Its ability to spread antibiotic resistance-carrying broad host range plasmids like P3 renders *S*. Tm a concerning driver of antibiotic resistance spread on a global scale.

The experimental setup comprised some limitations as we could not distinguish between transconjugants resulting from the initial donor and those resulting from secondary conjugation by transconjugants that were just replicating in the gut lumen. Thus, no precise conclusions about the actual transfer dynamics or *in vivo* plasmid loss can be drawn at this state. It is also still uncertain to what extent transconjugants serve as P3 donors *in vivo.* Nonetheless, our findings illustrate how antibiotic resistance genes can spread within the gut environment independently of selective pressure.

*S*. Tm might be of particular importance in driving conjugative HGT for two reasons. It can not only form tissue reservoirs including persisters (as discussed above), but also causes severe inflammation and thus leads to dysbiosis enabling blooms of potential recipient cells (esp. members of the Proteobacteria (53)) and thereby promote conjugation (27). In general, this further confirms how dysbiosis (which can be induced by many factors like diet, stress, antibiotic treatment or diseases (54–57)) might boost HGT and antibiotic resistance spread as soon as a potent donor strain like *S*. Tm SL1344 enters the gut system (27). As P2 is a close relative to the clinically relevant ESBL-plasmids (29), it might be interesting to investigate how well ESBL-plasmids are able to mobilize P3 or closely related plasmids from the IncQ family. Overall, the presented data add to our understanding of resistance plasmid transfer and suggest that conjugation and plasmid maintenance in gut luminal microbes and environmental bacteria should be considered in health approaches aiming to decipher and reduce the spread of antibiotic resistance.

## MATERIALS AND METHODS

### Bacterial strains and growth conditions

The strains and plasmids used in this study are listed in Table S1 and S2. All cells were routinely grown either on 1.5% Lysogeny Broth (LB) agar, MacConkey agar or in liquid LB supplemented with ampicillin (100 μg/ml), kanamycin (50 μg/ml), streptomycin (50 μg/ml) or chloramphenicol (35 μg/ml), where necessary. Gene deletions were obtained via PCR-based inactivation (58), and Km^r^ cassettes were eliminated via FLP recombination.

### Recipient screen

To study the host range of the P3 plasmid, bacterial strains from environmental and human isolates were screened for spectrum of antibiotic resistance and the ability to ferment lactose to enable later differentiation from the *S*. Tm donor strains on MacConkey agar plates.

To detect antibiotic resistances for all potential recipient strains, a minimal inhibition concentration (MIC) assay was conducted for ampicillin, streptomycin, chloramphenicol and kanamycin. 24 well plates (TPP, No. 92024) were filled with 1 mL Lysogeny Broth (LB) medium and supplemented with a range of concentrations of the respective antibiotics reaching from 0 – 100 μg/mL. Tested strains were grown in 5 mL LB overnight at 30 °C or 37 °C. Optical density (OD_600_) of the cultures was adjusted to 1.0. Subsequently, 10 μL of the culture was added to each well. The plates were incubated at 30 °C or 37 °C overnight and OD_600_ was measured for each well.

### *In vitro* mating

2 h liquid mating was performed to determine conjugation frequencies. The donor and recipient strains were grown individually and 10 μL (~ 1×10^7^ CFUs) of both, donor and recipient cells, were added to 1 mL LB. The cells were incubated at 37 °C for 2 h without shaking. To determine the number of donors, recipients and transconjugants, dilution series were plated on MacConkey or LB agar (depending on the recipient strain) supplemented with respective antibiotics. The conjugation frequency (CF) was determined by using formula [1].

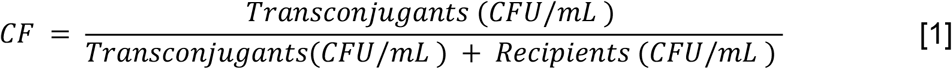

To determine whether P3 was expressed stably in a recipient bacterium, the same experiment was conducted for a longer period. 20 μL of the 1:1 mix of donor and recipient cells were added to 5 mL of LB medium without antibiotics and incubated at overnight without shaking. The numbers of donors, recipients and transconjugants were determined by respective plating. The ratio between transconjugant counts and recipient counts (ratioT/R) was used to normalize the dataset in order to compare different recipient strains [2].

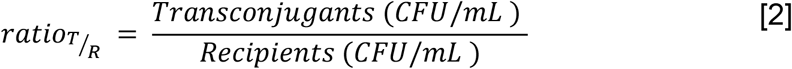

Mating experiments on agar as a solid surface were added to this study as conjugation events requires close physical contact of recipient and donor cells (5). Agar mating enabled an additional way of investigating the host range of P3 because it allowed higher densities of recipient and donor strains at the same spot than liquid mating. The donor and recipient strains were cultured and prepared as described above and 10 μL droplets of a 1:1 mix were pipetted onto LB agar without antibiotics. After incubating overnight, the cultures were scraped off the agar and resuspended in 1 mL of 1X PBS. Donors, recipients and transconjugants were enumerated as described above.

### Isolation of human Enterobacteriaceae from stool samples

The feces originated from healthy donors and from Salmonellosis patients in remission of two different clinical trials. The feces were collected and homogenized with tryptic soy broth + glycerol before storing at −80°C. For plating, an aliquot of the sample was scraped off the frozen stock and homogenized for 1 min at 25 Hz in PBS using a tissue lyzer (QIAGEN). Serial dilutions were prepared in PBS and plated on MacConkey Agar (OXOID, CM0007; incubation over-night at 37°C under aerobic conditions). Morphologically different colonies were picked and streaked on MacConkey to ensure purity of the culture (incubation over-night at 37°C under aerobic conditions). A single colony was picked to inoculate a 3 mL liquid LB culture (37°C, overnight). The pellet of the culture was used for DNA extraction (QIAGEN DNeasy Blood & Tissue Kit).

### DNA extraction and sequencing

The QIAGEN DNeasy Blood & Tissue Kit protocol was used and the DNA was eluted in 50 μL elution buffer (AE). The DNA samples were sent to Novogene for library preparation, sequencing (Illumina NovaSeq 6000) and raw read filtering (BioProject Accession Number PRJNA853708).

### Bioinformatics

#### Identification of the species

The paired-end reads were de-novo assembled with CLC Workbench 20.0.4. Contigs with lengths below 500 pb were discarded and the species of the isolates was identified by multi-locus sequence typing (59). The assemblies of the leaf isolates (47) (Fig S1) were downloaded from GenBank® (NCBI, BioProject Accession Numbers PRJNA224116 and PRJNA297956).

#### Search for repZ

The sequence of the *repZ* gene was downloaded from NCBI extracted from *S.* Tm SL1344 (Accession Number HE654725.1, (17)) and used for two sorts of BLAST® (60) searches (megablast and blastn) on the contigs/assemblies.

### Mouse models

Short term experiments (6 days) were carried out with 8–12-week-old, specific pathogen-free (SPF) C57BL/6 mice (JAX:000664, The Jackson Laboratory). Long term experiments (9-15 days) were carried out with 8–12-week-old SPF 129S6/SvEvTac mice (RRID:IMSR_TAC:129sve). All mice were bred at the EPIC mouse facility at ETH Zürich. During the experiment, they were housed under barrier conditions in individually ventilated cages at the ETH Zürich rodent center (RCHCI) with a maximum of 5 mice per cage. The mice were fed with a mouse maintenance diet (Kliba Nafag, 3537; autoclaved; per weight: 4.5% fat, 18.5% protein, ~50% carbohydrates, 4.5% fiber). Mice of both sexes were randomly assigned to experimental groups. All infection experiments were approved by the responsible authorities (Tierversuchskommission Kantonales Veterinäramt Zürich, license ZH158/2019).

#### Model for interspecific conjugation

Ampicillin-pretreated 8–12-week-old SPF C57BL/6 mice were orally infected by gavaging 5 x 10^7^ CFUs of the recipient strain one day prior to oral infection with 5 x 10^7^ CFUs of the donor strain. For preparing the inoculum, the relevant strains were cultured in LB with selective antibiotics overnight and subcultured (1:20 dilution) until they reached an OD_600_ of 0.5-1.0 (exp. growth phase) and adjusted to OD 1.0 by washing and diluting in sterile 1X PBS. The transfer of P3 was monitored for a total of 3 days by respective plating of fecal samples for donor, recipient and transconjugant populations on MacConkey or LB agar (depending on the recipient strain). Fecal samples were collected every 24 hours and placed in pre-weighed (wb) 2 mL Eppendorf tubes which contained 500 μL of sterile 1X PBS and a metal bead. The tubes were weighed again (wa) after sample collection and the samples were homogenized for 2 min at 25 Hz using a TissueLyser from Qiagen. At 4 d.p.i, the mice were sacrificed by CO_2_ asphyxiation. Cecal content as well as mesenteric lymph nodes, liver and spleen were collected and plated. The number of cells per organ was calculated by multiplying the counted CFUs by a factor of 10 for lymph nodes and spleen and by a factor of 60 for the liver, as only 1/6 of it was collected and plated. The formula [3] was used to determine the number of CFUs per gram feces or cecal content.

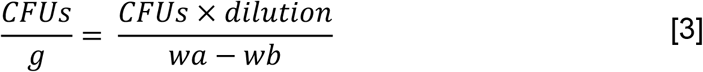

#### Reseeding donor model

*S*. Tm can invade host cells during a gastrointestinal infection and colonize systemic organs. Its ability to pass the epithelial barrier within the gut system by employing a type III secretion system can lead to reseeding events into the gut lumen (23, 61, 62). The described model was adapted from previous studies (23, 29). Ampicillin-pretreated 8–12-week-old SPF 129S6/SvEvTac mice were chosen to investigate the intraspecific transfer of P3 by a reseeding *S*. Tm donor for a total of 8 days. In contrast to C57BL/6, this mouse line has a functional allele for Nramp1 (also known as Slc11a1). The encoded solute carrier protein is a host resistance factor and allows for long term infections with *S*. Tm as it restricts *Salmonella* growth through deprivation of the crucial micronutrient Mn^2+^ (63, 64). The experiment contained two phases: 1.) Day 0 - 2: donor colonization and 2.) Day 2 - 8: recipient colonization, donor reseeding and plasmid transfer. 1.) Mice were i.p. infected with 10^3^ CFUs of the donor strain *S.* Tm SL1344 WITS1-KmR pM965 on day 0 (subcultured for 2.5 h and OD_600_ adjusted to 1.0). 2.) Mice were pretreated with 20 mg ampicillin on day 2 and 4 h later infected by oral gavage with 5 x 10^7^ CFUs of the recipient strain *S*. Tm 14028S *marT::cat* pM965 (subcultured for 2.5 h and OD_600_-adjusted as described above). Sterile filtered ampicillin (2 g/L) was added to the drinking water on day 2 after infection and maintained throughout the experiment to ensure continuous suppression of microbiota growth and stable colonization of donor and recipient strains. The transfer of P3 was monitored for a total of 5 days (day 4 – 8) by respective plating of fecal samples (day 4 - 7) and cecal content, mesenteric lymph nodes (mLN), liver and spleen (day 8) for donor, recipient and transconjugant cells. The mice were housed in individual cages throughout the experiment to prevent cross-contamination and minimize the risk of donor infections via the fecal-oral route and were euthanized by CO_2_ asphyxiation 8 d.p.i. The entire experimental setup is illustrated in Fig 3a, b.

#### Persisting donor model

The experiment consisted of three phases, 1.) Day 0 - 2: donor colonization. 2.) Day 2 – 7: antibiotic clearance 3.) Day 7 – 14: recipient colonization, reseeding of the persisting donors from tissue reservoirs, conjugative plasmid transfer. 1.) Mice were i.p infected with the donor (10^3^ CFUs) on day 0. 2.) On day 3, 4 and 5 the mice were treated with 1.5 mg ceftriaxone intraperitoneally to induce persister formation of the donor strain. The mice were put in separate cages after the last ceftriaxone treatment to minimize donor infections via coprophagy and cross-contamination. Additionally, ampicillin (2 g/L) was added to the drinking water on day 4 and maintained throughout the experiment to suppress the microbiota for a later introduction of the recipient strain *S*. Tm 14028S *marT::cat* pM965. The recipient was introduced orally on day 7 by gavaging 5 x 10^7^ CFUs. The inoculum was prepared as mentioned above. Feces were collected and plated on MacConkey with respective antibiotics on day 0, day 2 and day 7 to check for (unwanted) donor colonization in the gut and on day 9 – 14 to enumerate donor, recipient and transconjugant populations. The mice were euthanized at 14 d.p.i. by CO_2_ asphyxiation and cecal content, mNL, liver and spleen were collected and plated as described above. The experimental setup can be looked up in Fig 3 d, e.

### Data analysis

Microsoft Excel 2016 for Windows was used to calculate the cells/g content for all *in vivo* experiments. Data was analyzed and plotted using GraphPad Prism version 9.2.0 for Windows (GraphPad Software, La Jolla California USA, www.graphpad.com). For statistical analysis, the significance level was set to 5 %.

## DATA AVAILABILITY

All raw data and executed protocols gathered during this study are available upon request from Prof. Dr. Wolf-Dietrich Hardt.

## CONTRIBUTIONS

M.G., L.L., W.-D.H. and E.B. conceived and designed the experiments, M.G. and L.L. performed the experiments. M.H. performed bioinformatic analysis. All authors contributed to data analysis and writing of the manuscript.

## ACKNOWLEDGMENTS

We would like to thank the members of the Hardt lab for fruitful discussion and acknowledge excellent support from the EPIC mouse facility of ETH Zurich, in particular Sanne Kroon, Yves Steiger, Noemi Santamaria de Souza, Ersin Gül, Prof. Dr. Julia Vorholt and Dr. Martin Schäfer for providing help and strains. This work has been supported by grants from the Swiss National Science Foundation (310030_192567 and NCCR Microbiomes) and the European Union’s Horizon 2020 Research and Innovation Program under the Marie Skłodowska-Curie grant agreement 956279 COL_RES to WDH. L.L. is supported by LA 4572/1-1 grant from the Deutsche Forschungsgemeinschaft.

## REFERENCES

1. Clatworthy AE, Pierson E, Hung DT. Targeting virulence: a new paradigm for antimicrobial therapy. Nat Chem Biol. 2007;3(9):541–8.

2. Van Boeckel TP, Brower C, Gilbert M, Grenfell BT, Levin SA, Robinson TP, et al. Global trends in antimicrobial use in food animals. Proc Natl Acad Sci U S A. 2015;112(18):5649–54.

3. McManus PS, Stockwell VO, Sundin GW, Jones AL. Antibiotic use in plant agriculture. Annu Rev Phytopathol. 2002;40:443–65.

4. WHO. WHO publishes list of bacteria for which new antibiotics are urgently needed: World Health Organization, Media Centre; 2017 [Available from: https://www.who.int/news/item/27-02-2017-who-publishes-list-of-bacteria-for-which-new-antibiotics-are-urgently-needed.

5. Murray CJL, Ikuta KS, Sharara F, Swetschinski L, Robles Aguilar G, Gray A, et al. Global burden of bacterial antimicrobial resistance in 2019: a systematic analysis. The Lancet. 2022;399(10325):629–55.

6. Stockwell VO, Duffy B. Use of antibiotics in plant agriculture. Rev sci tech Off int Epiz. 2012;1(31):199–210.

7. He Y, Yuan Q, Mathieu J, Stadler L, Senehi N, Sun R, et al. Antibiotic resistance genes from livestock waste: occurrence, dissemination, and treatment. npj Clean Water. 2020;3(1).

8. Ochman H, Lawrence JG, Groisman EA. Lateral gene transfer and the nature of bacterial innovation. Nature. 2000;405:299–304.

9. Brito IL. Examining horizontal gene transfer in microbial communities. Nat Rev Microbiol. 2021;19(7):442–53.

10. Virolle C, Goldlust K, Djermoun S, Bigot S, Lesterlin C. Plasmid transfer by conjugation in Gram-negative bacteria: from the cellular to the community level. Genes (Basel). 2020;11(11):1239.

11. Norman A, Hansen LH, Sorensen SJ. Conjugative plasmids: vessels of the communal gene pool. Philos Trans R Soc Lond B Biol Sci. 2009;364(1527):2275–89.

12. von Wintersdorff CJ, Penders J, van Niekerk JM, Mills ND, Majumder S, van Alphen LB, et al. Dissemination of antimicrobial resistance in microbial ecosystems through horizontal gene transfer. Front Microbiol. 2016;7:173.

13. Summers DK. The Biology of plasmids: Blackwell Science Ltd., Oxford; 1996.

14. Eberhard WG. Why do bacterial plasmids carry some genes and not others? Plasmid. 1989;21:167–74.

15. Smillie C, Garcillan-Barcia MP, Francia MV, Rocha EP, de la Cruz F. Mobility of plasmids. Microbiol Mol Biol Rev. 2010;74(3):434–52.

16. Kingsley RA, Bäumler J. Host adaptation and the emergence of infectious disease - the Salmonella paradigm. Mol Microbiol. 2000;36(5):1006–14.

17. Kroger C, Dillon SC, Cameron AD, Papenfort K, Sivasankaran SK, Hokamp K, et al. The transcriptional landscape and small RNAs of *Salmonella enterica* serovar Typhimurium. Proc Natl Acad Sci U S A. 2012;109(20):E1277–E86.

18. Rankin JD, Taylor RJ. The estimation of doses of *Salmonella* Typhimurium suitable for the experimental production of disease in calves.. Vet Rec. 1966;78(21):706–7.

19. Rawlings DE, Tietze E. Comparative biology of IncQ and IncQ-like plasmids. Microbiol Mol Biol Rev. 2001;65(4):481–96.

20. Guerry P, van Embden J, Falkow S. Molecular nature of two nonconjugative plasmids carrying drug resistance genes. J Bacteriol. 1974;117(2):619–30.

21. Yau S, Liu X, Djordjevic SP, Hall RM. RSF1010-like plasmids in Australian *Salmonella enterica* serovar Typhimurium and origin of their *sul2-strA-strB* antibiotic resistance gene cluster. Microb Drug Resist. 2010;16(4):249–52.

22. Tietze E, Tschäpe H, Voigt W. Characterization of new resistance plasmids belonging to incompatibility group *IncQ* J Basic Microbiol. 1989;29(10):695–706.

23. Bakkeren E, Huisman JS, Fattinger SA, Hausmann A, Furter M, Egli A, et al. *Salmonella* persisters promote the spread of antibiotic resistance plasmids in the gut. Nature. 2019;573(7773):276–80.

24. Sanchez-Romero MA, Merida-Floriano A, Casadesus J. Copy number heterogeneity in the virulence plasmid of *Salmonella enterica*. Front Microbiol. 2020;11:599931.

25. Asano K, Mizobuchi K. Copy number control of IncIα plasmid ColIb-P9 by competition between pseudoknot formation andantisense RNA binding at a specific RNA site. EMBO. 1998;17(17):5201–13.

26. Hiley L, Graham RMA, Jennison AV. Genetic characterisation of variants of the virulence plasmid, pSLT, in *Salmonella enterica* serovar Typhimurium provides evidence of a variety of evolutionary directions consistent with vertical rather than horizontal transmission. PLoS One. 2019;14(4):e0215207.

27. Stecher B, Denzler R, Maier L, Bernet F, Sanders MJ, Pickard DJ, et al. Gut inflammation can boost horizontal gene transfer between pathogenic and commensal Enterobacteriaceae. Proc Natl Acad Sci U S A. 2012;109(4):1269–74.

28. Scherzinger E, Bagdasarian MM, Scholz P, Lurz R, Rückert B, Bagdasarian M. Replication of the broad host range plasmid RSF1010-requirement for three plasmid-encoded proteins. PNAS. 1983;81:654–8.

29. Bakkeren E, Herter JA, Huisman JS, Steiger Y, Gul E, Newson JPM, et al. Pathogen invasion-dependent tissue reservoirs and plasmid-encoded antibiotic degradation boost plasmid spread in the gut. Elife. 2021;10:e69744.

30. Nair DVT, Venkitanarayanan K, Kollanoor Johny A. Antibiotic-resistant *Salmonella* in the food supply and the potential role of antibiotic alternatives for control. Foods. 2018;7(10).

31. Naylor NR, Atun R, Zhu N, Kulasabanathan K, Silva S, Chatterjee A, et al. Estimating the burden of antimicrobial resistance: a systematic literature review. Antimicrob Resist Infect Control. 2018;7:58.

32. Jajere SM. A review of *Salmonella enterica* with particular focus on the pathogenicity and virulence factors, host specificity and antimicrobial resistance including multidrug resistance. Vet World. 2019;12(4):504–21.

33. Gormey EP, Davies J. Transfer of plasmid RSF1010 by conjugation from *Escherichia coli* to *Streptomyces lividans* and *Mycobacterium smegmatis*. J Bacteriol. 1991;173(21):6705–8.

34. De Gelder L, Ponciano JM, Joyce P, Top EM. Stability of a promiscuous plasmid in different hosts: no guarantee for a long-term relationship. Microbiology (Reading). 2007;153(Pt 2):452–63.

35. Foley SL, Kaldhone PR, Ricke SC, Han J. Incompatibility Group I1 (IncI1) plasmids: Their genetics, biology, and public health relevance. Microbiol Mol Biol Rev. 2021;85(2).

36. Barthel M, Hapfelmeier S, Quintanilla-Martinez L, Kremer M, Rohde M, Hogardt M, et al. Pretreatment of mice with streptomycin provides a *Salmonella enterica* serovar Typhimurium colitis model that allows analysis of both pathogen and host. Infect Immun. 2003;71(5):2839–58.

37. Monack DM, Mueller A, Falkow S. Persistent bacterial infections: the interface of the pathogen and the host immune system. Nat Rev Microbiol. 2004;2(9):747–65.

38. Lawley TD, Chan K, Thompson LJ, Kim CC, Govoni GR, Monack DM. Genome-wide screen for *Salmonella* genes required for long-term systemic infection of the mouse. PLoS Pathog. 2006;2(2):e11.

39. Kaiser P, Regoes RR, Dolowschiak T, Wotzka SY, Lengefeld J, Slack E, et al. Cecum lymph node dendritic cells harbor slow-growing bacteria phenotypically tolerant to antibiotic treatment. PLoS Biol. 2014;12(2):e1001793.

40. Balaban NQ, Merrin J, Chait R, Kowalik L, Leibler S. Bacterial Persistence as a Phenotypic Switch. Science. 2004;305:1622–5.

41. Manuse S, Shan Y, Canas-Duarte SJ, Bakshi S, Sun WS, Mori H, et al. Bacterial persisters are a stochastically formed subpopulation of low-energy cells. PLoS Biol. 2021;19(4):e3001194.

42. Wakamoto Y, Dhar N, Chait R, Schneider K, Signorino-Gelo F, Leibler S, et al. Dynamic persistence of antibiotic-stressed Mycobacteria. Science. 2013;339:91–5.

43. Brauner A, Fridman O, Gefen O, Balaban NQ. Distinguishing between resistance, tolerance and persistence to antibiotic treatment. Nat Rev Microbiol. 2016;14(5):320–30.

44. Claudi B, Sprote P, Chirkova A, Personnic N, Zankl J, Schurmann N, et al. Phenotypic variation of *Salmonella* in host tissues delays eradication by antimicrobial chemotherapy. Cell. 2014;158(4):722–33.

45. Fisher RA, Gollan B, Helaine S. Persistent bacterial infections and persister cells. Nat Rev Microbiol. 2017;15(8):453–64.

46. Garcia-Quintanilla M, Casadesus J. Virulence plasmid interchange between strains ATCC 14028, LT2, and SL1344 of *Salmonella enterica* serovar Typhimurium. Plasmid. 2011;65(2):169–75.

47. Bai Y, Muller DB, Srinivas G, Garrido-Oter R, Potthoff E, Rott M, et al. Functional overlap of the *Arabidopsis* leaf and root microbiota. Nature. 2015;528(7582):364–9.

48. Sundin GW, Bender CL. Dissemination of the *strA-strB* streptomycin-resistance genes among commensal and pathogenic bacteria from humans, animals, and plants. Mol Ecology. 1996(5):133–43.

49. van Overbeek LS, Wellington EMH, Egan S, Smalla K, Heuer H, Collard J-M, et al. Prevalence of streptomycin-resistance genes in bacterial populations in European habitats. FEMS Microbiol Ecol. 2002;42:277–88.

50. Bahl MI, Hansen LH, Sorensen SJ. Impact of conjugal transfer on the stability of IncP-1 plasmid pKJK5 in bacterial populations. FEMS Microbiol Lett. 2007;266(2):250–6.

51. Bergstrom CT, Lipsitch M, Levin BR. Natural Selection, Infectious Transfer and the existence conditions for bacterial plasmids. Genetics. 2000;155(4):1505–19.

52. Okoro CK, Kingsley RA, Quail MA, Kankwatira AM, Feasey NA, Parkhill J, et al. High-resolution single nucleotide polymorphism analysis distinguishes recrudescence and reinfection in recurrent invasive nontyphoidal *Salmonella* Typhimurium disease. Clin Infect Dis. 2012;54(7):955–63.

53. Spees AM, Lopez CA, Kingsbury DD, Winter SE, Baumler AJ. Colonization resistance: battle of the bugs or Menage a Trois with the host? PLoS Pathog. 2013;9(11):e1003730.

54. Brown K, DeCoffe D, Molcan E, Gibson DL. Diet-induced dysbiosis of the intestinal microbiota and the effects on immunity and disease. Nutrients. 2012;4(8):1095–119.

55. De Palma G, Collins SM, Bercik P, Verdu EF. The microbiota-gut-brain axis in gastrointestinal disorders: stressed bugs, stressed brain or both? J Physiol. 2014;592(14):2989–97.

56. Francino MP. Antibiotics and the human gut microbiome: dysbioses and accumulation of resistances. Frontiers in Microbiology. 2016;6.

57. Wilkins LJ, Monga M, Miller AW. Defining dysbiosis for a cluster of chronic diseases. Sci Rep. 2019;9(1):12918.

58. Datsenko KA, Wanner BL. One-step inactivation of chromosomal genes in *Escherichia coli* K-12 using PCR products. Proceedings of the National Academy of Sciences. 2000;97(12):6640–5.

59. Jolley KA, Bray JE, Maiden MCJ. Open-access bacterial population genomics: BIGSdb software, the PubMLST.org website and their applications. Wellcome Open Res. 2018;3:124.

60. Zhang Z, Schwartz S, Wagner L, Miller W. A Greedy Algorithm for aligning DNA sequences. Journal of Computational Biology. 2000;7:203–14.

61. Waterman SR, Holden DW. Functions and effectors of the *Salmonella* pathogenicity island 2 type III secretion system. Cellular Microbiology. 2003;5(8):501–11.

62. Lam LH, Monack DM. Intraspecies competition for niches in the distal gut dictate transmission during persistent *Salmonella* infection. PLoS Pathog. 2014;10(12):e1004527.

63. Stecher B, Paesold G, Barthel M, Kremer M, Jantsch J, Stallmach T, et al. Chronic *Salmonella enterica* serovar Typhimurium-induced colitis and cholangitis in streptomycin-pretreated Nramp1+/+ mice. Infect Immun. 2006;74(9):5047–57.

64. Cunrath O, Burmann D. Host resistance factor SLC11A1 restricts *Salmonella* growth through magnesium deprivation. Science. 2019(366):995–9.

